# Towards a Neurometric-based Construct Validity of Trust

**DOI:** 10.1101/2021.07.04.451074

**Authors:** Pin-Hao A. Chen, Dominic Fareri, Berna Güroğlu, Mauricio R. Delgado, Luke J. Chang

**Author notes:** Corresponding author &.

## Abstract

Trust is a nebulous construct central to successful cooperative exchanges and interpersonal relationships. In this study, we introduce a new approach to establishing construct validity of trust using “neurometrics”. We develop a whole-brain multivariate pattern capable of classifying whether new participants will trust a relationship partner in the context of a cooperative interpersonal investment game (n=40) with 90% accuracy and find that it also generalizes to a variant of the same task collected in a different country with 82% accuracy (n=17). Moreover, we establish the convergent and discriminant validity by testing the pattern on thirteen separate datasets (n=1,427) and find that trust is reliably related to beliefs of safety, inversely related to negative affect, but unrelated to reward, cognitive control, social perception, theory of mind, and self-referential processing. Together these results provide support for the notion that the psychological experience of trust contains elements of beliefs of reciprocation and fear of betrayal aversion. Contrary to our predictions, we found no evidence that trust is related to anticipated reward. This work demonstrates how “neurometrics” can be used to characterize the psychological processes associated with brain-based multivariate representations.

## Introduction

The foundation of modern society is built upon our ability to successfully conduct cooperative social exchanges such as strategic coalitions, exchange markets, and systems of governance. Trust plays a central role in facilitating social exchange (Arrow, 1974) based on its ability to reduce transaction costs and increase information sharing (Dyer and Chu, 2003). Successful interpersonal, business, and political transactions require trusting that a relationship partner will honor their agreement. Countries with formal institutions that protect property and contract rights are associated with higher perceptions of trust and civic cooperation, decreased rates of violent crime in neighborhoods (Sampson et al., 1997), and increased economic growth (Knack and Keefer, 1997). From an interpersonal perspective, trust can be considered the psychological state of assuming mutual risk with a relationship partner to attain an interdependent goal in the face of competing temptations *(Chang, 2017; Simpson, 2007)* which can be assayed using a two-person Investment Game (Berg et al., 1995; Dufwenberg and Gneezy, 2000). In this game, a Trustor has the opportunity to invest a portion of a financial endowment to a Trustee. The investment amount is multiplied by a factor specified by the experimenter (e.g., 3 or 4), and the Trustee ultimately decides how much of the multiplied endowment to return back to the Trustor to honor or betray their trust. This game has been well studied in behavioral economics (Camerer, 2003) and also in the field of decision neuroscience, which has investigated the neurobiological processes associated with trust (Delgado et al., 2005; Fareri et al., 2015, 2012a; Fouragnan et al., 2013; King-Casas et al., 2008, 2005; Krajbich et al., 2009; McCabe et al., 2001; Phan et al., 2010) and its reciprocation (Chang et al., 2011; van Baar et al., 2019). This work has found that trust and reciprocity are associated with neural reward circuitry including the ventral striatum, ventral tegmental area (VTA), and medial prefrontal cortex. However, it remains unclear precisely how this neural circuitry produces psychological feelings of trust that drives behavior in interpersonal interactions. In this paper, we establish a “neurometric” approach to assess the construct validity of brain activity patterns predictive of individual decisions to trust in the investment game.

Trust is a dynamic state that evolves over the course of a relationship. Early stages of a relationship are focused on assessing a partner’s trustworthiness level, which can be impacted by previous interactions (Delgado et al., 2005; Fareri et al., 2012a; Fouragnan et al., 2013; Frank et al., 1993), gossip (Feinberg et al., 2014; Jolly and Chang, 2018; Sommerfeld et al., 2007), group membership (Stanley et al., 2011), or judgments based on appearance (Scharlemann et al., 2001; van ’t Wout and Sanfey, 2008). Trustors must be willing to endure a risk (Rousseau et al., 1998; Scanzoni, 1979), while Trustees must be willing to overcome their own self-interest and take an action that fulfills an interdependent goal. As the relationship progresses, both parties are better able to predict each other’s behavior and develop a sense of security in the relationship. Trustors are more likely to trust a relationship partner when they have greater certainty that their partner will reciprocate (Chang et al., 2011). In this way, trustworthiness reflects a dynamic belief about the likelihood of a relationship partner reciprocating (Chang et al., 2010; Fareri et al., 2012a; King-Casas et al., 2005; Kishida and Montague, 2012; Phan et al., 2010; Rilling et al., 2002). These mutually beneficial collaborations can be rewarding (Delgado et al., 2005; Fareri et al., 2015; Phan et al., 2010; Tabibnia et al., 2008). However, at some point in the relationship, one person may end up betraying their partner (King-Casas et al., 2008), which could eventually lead to a dissolution of the relationship (King-Casas et al., 2008). Even the thought of being betrayed can be aversive enough to reduce trust behavior (Aimone et al., 2014; Aimone and Houser, 2012; Bohnet and Zeckhauser, 2004). Though both players may care about the others’ outcome, which requires reasoning about others’ mental states (McCabe et al., 2001) and concern for others’ welfare (Cox, 2004), there is little evidence behaviorally that trust is specifically motivated by altruism (Bohnet and Zeckhauser, 2004; Brülhart and Usunier, 2012). Thus, the candidate motivations influencing our likelihood to place trust in others include: (a) beliefs about probability of future reciprocation, (b) anticipated rewards, and (c) betrayal-aversion.

In psychometrics, creating a quantitative measurement of a nebulous and multifaceted concept such as trust requires establishing construct validity. Constructs provide consensus understanding of the semantic meaning of an abstract concept based on a nomological network of associations to other concepts (Cronbach and Meehl, 1955). Validating a construct requires assessing its generalizability to new populations and contexts and its convergent and discriminant validity to other constructs (Campbell and Fiske, 1959). Although the principles of psychometrics were originally established for more traditional psychological tests and questionnaires, there is growing evidence that patterns of brain activity can serve as “neurometrics” of constructs (Woo et al., 2017). For example, there has been a longstanding interest in using functional localizers (Saxe et al., 2006) and multivariate decoding methods to determine an individual’s psychological state based on patterns of brain activity (Chang et al., 2013; Kragel et al., 2018; Poldrack et al., 2009; Yarkoni et al., 2011) with demonstrated success in predicting the intensity of a variety of affective experiences (Chang et al., 2015; Eisenbarth et al., 2016; Krishnan et al., 2016; Wager et al., 2013; Yu et al., 2020), reconstructing a visual stimulus (Nishimoto et al., 2011) or uncovering its semantic meaning (Huth et al., 2016). Neurometrics has several advantages over psychometrics in that it can utilize a high dimensional measurement of voxel activity observed during the engagement of a specific psychological process without requiring retrospective verbal self-report (e.g., questionnaires) or completing many different behavioral tasks (e.g., intelligence tests). By leveraging quickly changing scientific norms in open data sharing (Gorgolewski et al., 2017, 2016, 2015; Poldrack and Gorgolewski, 2014; Yarkoni et al., 2010), it is increasingly possible to train a model predictive of a psychological state using brain activity such as pain (Wager et al., 2013), and establish a nomological network based on the model’s convergent and discriminative validity with other constructs such as negative emotions (Chang et al., 2015), cognitive control (Kragel et al., 2018), social rejection (Woo et al., 2014), and vicariously experienced pain (Krishnan et al., 2016).

Building on this approach, in this study we use supervised multivariate pattern-based analysis to predict individual decisions to trust a relationship partner in an interpersonal context using data from two previously published studies (Fareri et al., 2015, 2012a) (Figure 1A). We then establish the neurometric properties of this brain model by assessing its generalizability to a slightly different version of the task collected in a different country (Schreuders et al., 2018) (Figure 1B) and its convergent and divergent validity across 13 different tasks probing risk (Poldrack et al., 2016; Schonberg et al., 2012), affect (Chang et al., 2015; Hsiao et al., 2023), rewards (Barch et al., 2013; Fareri et al., 2012b; Tomova et al., 2020; Zhang et al., 2017), cognitive control (Xue et al., 2008), and social cognition (Barch et al., 2013; Chen et al., 2019, 2015, 2013; Wakeman and Henson, 2015). This process allows us to characterize the psychological properties of the construct of trust using neurometric analyses (Figure 1C). We focus specifically on psychological constructs that can be reliably elicited by experimental paradigms in a neuroimaging environment, but note that these paradigms may lack subtle nuances of the constructs, which may be refined in future work as more tasks become available. Based on the findings outlined above, we hypothesize that the construct of trust will be positively associated with beliefs of safety, feelings of anticipated reward, and negatively with feelings of negative affect, but not associated with other psychological processes (Figure 1D).

**Figure 1.**
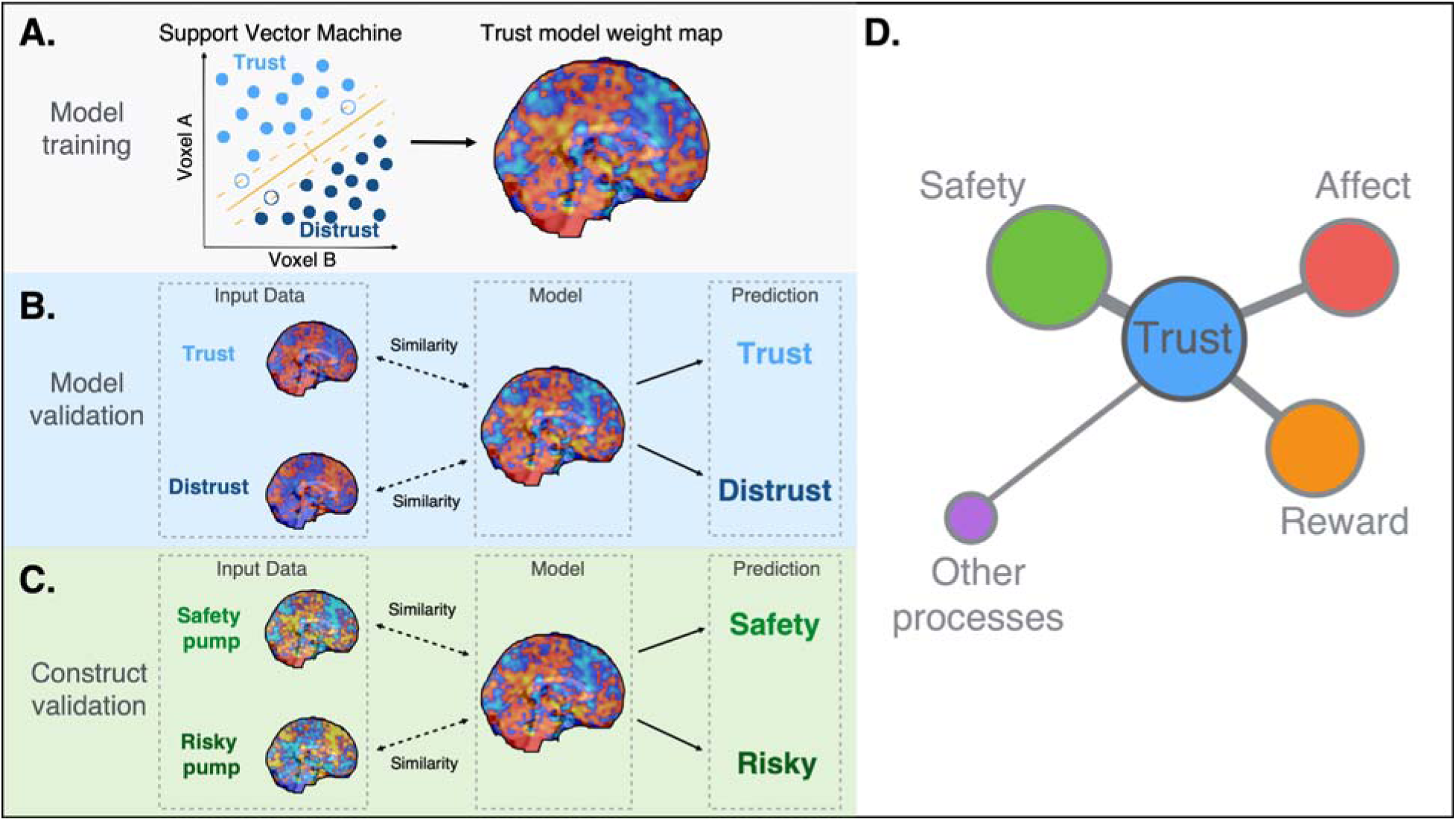
A demonstration of construct validity based on neurometric information. (A) A support vector machine algorithm was used to train the trust model. (B) An independent trust dataset was used to validate the trust model’s generalizability. (C) We tested the model on independent datasets such as the Balloon Analog Risk Task to assess the convergent and discriminant validity of the trust model. (D) We hypothesized that trust was associated with beliefs of safety, feelings of anticipated reward, and affect, but not other processes. The distances and thickness of edges represent the generalizability of the trust model to other domains, and the size of nodes represents sample size of the dataset from each domain.

## Results

### Training trust brain model

To establish a multivariate brain model of trust, we trained a linear Support Vector Machine (SVM) to classify when participants (n=40) decided to trust a relationship partner in the investment game using whole-brain patterns of brain activity measured in two previously published studies (Fareri et al., 2015, 2012a) (Figure 1A). The neural activity associated with taking an action to trust a relationship partner likely reflects objective signals of the psychological processes associated with the nebulous construct of trust. We performed an initial temporal data reduction using univariate general linear models (GLMs) to create an average map of each participant’s brain response when making decisions to trust or not. We then used a leave-one-subject-out (LOSO) cross-validation procedure to evaluate the performance of our multivariate SVM model in classifying maps associated with each participant’s decisions to prospectively trust or distrust using data from the rest of the participants. Our trust brain model (Figure 2A) was able to accurately discriminate between trust and distrust decisions within each participant (forced-choice accuracy: 90%, *p* < 0.001, Figure 2B & 2C, Table S1). Forced choice tests compare the relative pattern expression of the model between brain maps within the same participant and are particularly well suited for fMRI because they do not require signals to be on the same scale across individuals or scanners (Wager et al., 2013).

**Figure 2.**
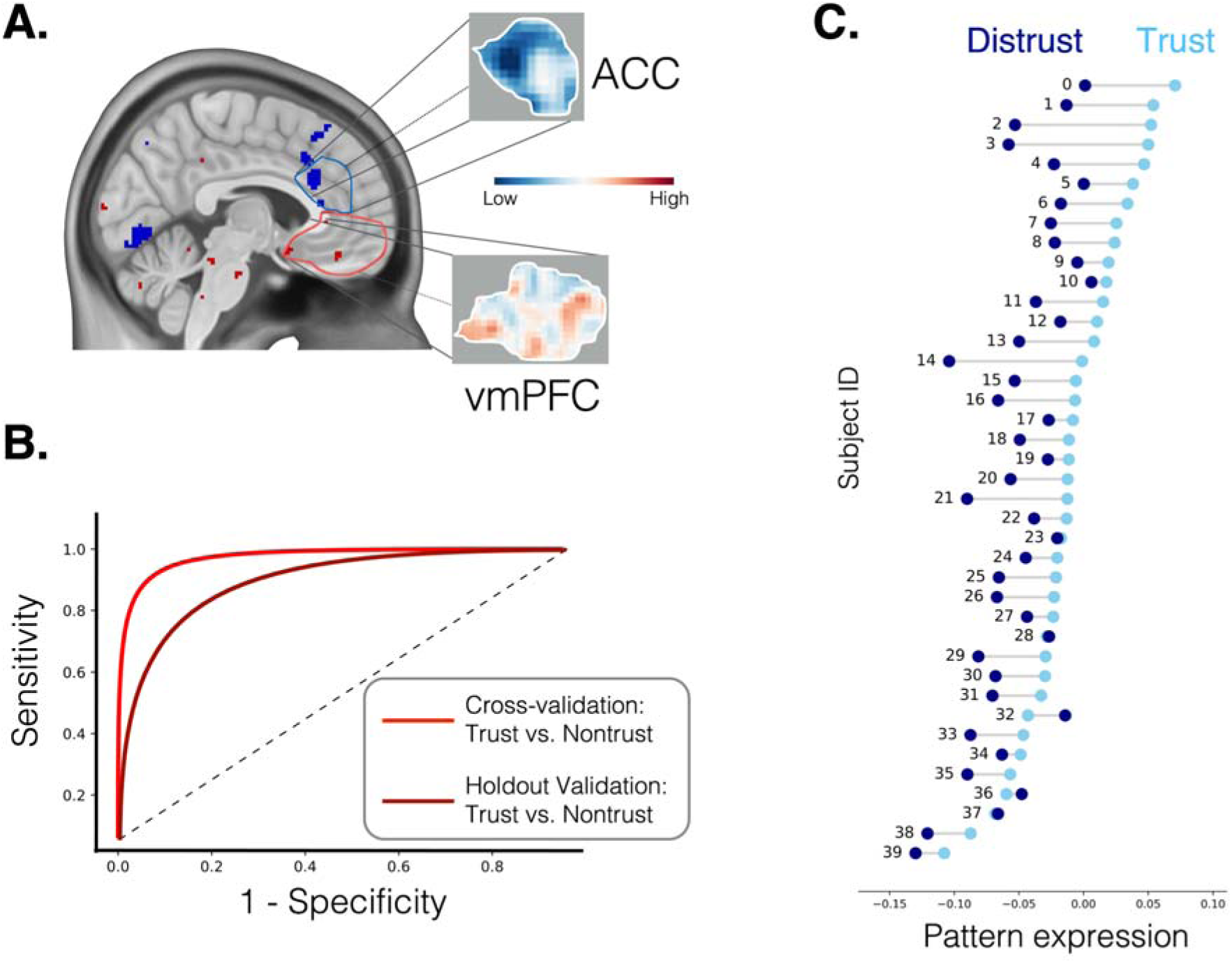
The trust model and its performance in the training and validation dataset. (A) The trust model is a whole brain pattern of voxel weights that can be linearly combined with new data to predict psychological levels of trust. We visualize the voxels that most reliably contribute to the classification using a bootstrap procedure (thresholded p < 0.005 uncorrected for visualization). (B) The receiver-operating-characteristic (ROC) plot highlights the sensitivity and specificity of the model in cross-validation and in an independent holdout dataset. (C) We plot the pattern expression, which is the spatial correlation between the cross-validated model and each participant’s average activity in the trust and distrust conditions based on their actual decisions in the game. Each participant (N=40) is depicted as a separate row with a line connecting the correlation values from each condition. The model correctly predicted the decisions of 36 out of 40 participants (90% accuracy).

To establish the face validity of our model, we used a parametric bootstrap to identify which voxels most reliably contributed to the classification, which involved retraining the model 5,000 times after randomly sampling participants with replacement. This procedure is purely for visualization and not used for spatial feature selection (Jolly and Chang, 2021). Consistent with prior work, we observed positive weights in the ventromedial prefrontal cortex (vmPFC), septal area (Fareri et al., 2015, 2012a; Krueger et al., 2007), amygdala, and ventral hippocampus. Negative weights were found in the dorsal anterior cingulate cortex (dACC) and bilateral insula (Figure 2A). These weights reflect how each voxel independently contributes to the classification of decisions to trust a relationship partner or not and were highly consistent. To estimate the reliability of the pattern, we performed a bootstrap analysis, in which we retrained the model after randomly sampling the data with replacement and computed the pairwise spatial similarity across each bootstrap iteration. Overall, we observed a high level of consistency of the spatial pattern, r=0.91 (Chang et al., 2015), which indicates that this pattern is likely to be robust to subtle variations in individual participant’s brain activity.

Next, we trained a general trust model using data from all participants and evaluated its generalizability on a variant of the trust game in which participants receive feedback about their partner’s decisions regardless if the participant decided to trust or not (Figure 1B). Importantly, we found that our model was able to accurately discriminate between the trust and distrust decisions from participants recruited from a different country collected on a different scanner (forced-choice accuracy: 82%, *p* = 0.006, Figure 2B, Table S1). This provides further confirmation that our model is capturing aspects of the psychological experience of trust that is shared across participants.

### Construct Validity

After establishing the sensitivity of our model to accurately discriminate trust decisions, we next sought to evaluate the generalizability of the trust model to other psychological constructs using additional datasets. If the model performs at chance in other contexts, then this establishes the specificity of the model in capturing trust. However, if the model gets confused in other contexts, then this may reflect overlap in the psychological experience of trust to other related constructs.

Trust reflects security in the relationship that the partner will behave as expected in their mutually interdependent interests (Chang et al., 2010). Beliefs about the likelihood of a particular outcome occurring (e.g., a relationship partner will reciprocate) can be represented as a probability distribution with certainty of safety at one end and risk of defection on the other. Risk and safety are thus inversely related. The entropy of this distribution can be described as uncertainty and reflects an inability to predict either a positive or negative outcome. We first examined whether the trust model might be related to beliefs of safety, which can be measured using risk-taking tasks. The Balloon Analog Risk task (BART) is among the most widely used behavioral assay of risk-taking behavior (Poldrack et al., 2016; Schonberg et al., 2012). In this task, participants are presented a series of colorful (the risk condition) or achromatic balloons (the safety or control condition) and are instructed to inflate the balloons. In the risk condition, participants can choose to inflate a balloon and only receive a reward if the balloon does not explode. However, each inflation is associated with an increasing probability of explosion, and when the balloon explodes, participants do not receive a reward for that round (Poldrack et al., 2016; Schonberg et al., 2012). In contrast, in the safety condition, participants are also instructed to inflate a series of balloons, but there is no risk of the balloons exploding, nor an opportunity to receive a reward. We calculated the spatial similarity of our trust model to univariate beta maps from a GLM measuring average brain activity to the risk or safety conditions from two independent BART datasets (N=15 in dataset 3(Schonberg et al., 2012) and N = 123 in dataset 4 (Poldrack et al., 2016); Table S1). In both datasets, we found that the trust model could accurately discriminate between the safety and risk conditions (accuracy=100%, *p* < 0.001 in dataset 3; accuracy=94%, *p* < 0.001 in dataset 4; Figure 3B-2). These results indicate that the trust model captures a psychological experience that is shared with beliefs about safety when making a decision indicating a low risk of a negative outcome occurring (Figure 3A-2). When a relationship partner seems untrustworthy and reciprocation seems risky, participants will choose to keep their money rather than investing it.

**Figure 3.**
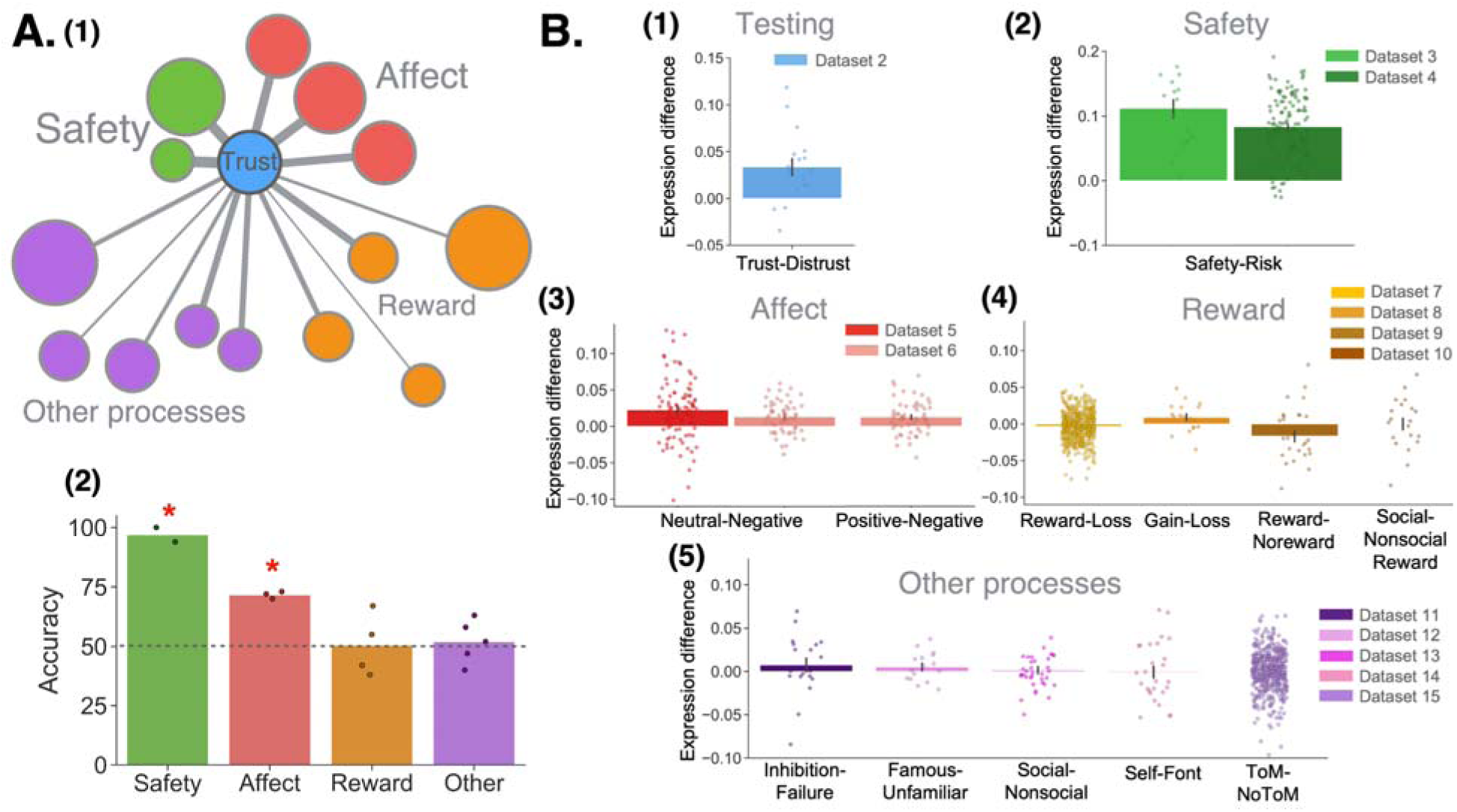
Construct validity of the trust model and model generalizability. (A) (1) Network plot illustrates that the trust model significantly generalizes to safety and affect datasets, but not to reward and other processing datasets. The distances and thickness of edges are weighted based on the rank of classification accuracy, and the size of nodes represents sample size of each dataset. (2) The forced-choice classification accuracy for each dataset within the four domains was shown in the bar plot. Only the safety and affect domains demonstrated above chance accuracy across datasets. (B) Trust model pattern expression differences between the two conditions in the: (1) trust testing datasets, (2) two safety datasets, (3) two affect datasets, (4) four reward datasets, as well as (5) five datasets involving cognitive control and social cognition.

Next, we explored if the trust model captured aspects of the experience related to negative affect. One reason why people may choose to distrust and not invest their money in a relationship partner is because of potential concerns about the partner betraying their trust and keeping all of the money. This results in negative utility for both losing money, and also being betrayed (Aimone et al., 2014; Bohnet and Zeckhauser, 2004). To test this hypothesis, we evaluated if the trust model might be inversely related to feelings of negative affect elicited by pictures from the international affective picture system (IAPS) from two independent datasets (Table S1). We found that in dataset 5 (N=93)(Chang et al., 2015), the trust model differentiated between conditions of neutral and negative emotional pictures (accuracy = 72%, *p* < 0.001; Figure 3B-3). A similar finding was also shown in dataset 6 (N=56) (Hsiao et al., 2023), where the trust model discriminated between the neutral and negative-valence picture conditions (accuracy = 70%, *p* = 0.002; Figure 3B-3) as well as between positive and negative-valence conditions (accuracy = 73%, *p* < 0.001; Figure 3B-3). These analyses provide evidence of overlap in the psychological processes associated with trust and negative affect. Specifically, decisions to trust are associated with less negative affect, consistent with a betrayal-aversion motivation. However, it is also possible that decisions to trust are associated with positive affect, but dataset 6 rules out this possibility as we did not observe a significant association with viewing positive compared to neutral pictures (accuracy = 50%, *p* = 0.553; Table S1), only positive and neutral compared to viewing negative pictures.

Third, we examined whether the trust model can be generalized to feelings of anticipated reward (Delgado et al., 2005; Phan et al., 2010). We have previously demonstrated that learning that a close friend reciprocated trust is associated with a greater rewarding experience compared to when a stranger reciprocates (Fareri et al., 2015), suggesting that trust may be associated with the anticipation of a future reward. To test this hypothesis, we evaluated whether our model was related to reward across four different tasks (Barch et al., 2013; Fareri et al., 2012b; Geier et al., 2010; Zhang et al., 2017). In dataset 7 (N=490; Table S1), participants played the Delgado Card Task and guessed whether a randomly drawn card would be higher or lower than a specific number. If they were correct, they would receive a monetary reward, and if they were incorrect they would lose money (Delgado et al., 2000). We found that the trust model performed at chance in differentiating experienced rewards from losses (Barch et al., 2013) (accuracy = 42%, *p* = 0.999; Figure 3B-4). Next we explored whether trust may be related to anticipated rewards. In dataset 8 (N=18; Table S1), participants were shown either a cue indicating maximal gain or loss (Zhang et al., 2017), which provides a way to measure the anticipation of a gain or loss. We found that the trust model was unable to discriminate between these two conditions (accuracy = 67%, *p* = 0.120; Figure 3B-4). Consistent with these results, the trust model was also unable to discriminate between anticipated rewards from no-rewards in the popular monetary incentive delay task (Knutson et al., 2000) in dataset 9 (N=29; Table S1; accuracy = 38%, *p* = 0.929; Figure 3B-4) (Tomova et al., 2020). Together, these studies elicit different aspects of reward from viewing a cue that predicts the future receipt of a reward (dataset 8), to the anticipation of receiving a reward (dataset 9), to the experience of receiving a reward (dataset 7) when the reward is a financial outcome. We were also interested in whether the trust model may capture social rewards, which would support the role of altruism or concern for the relationship partners’ outcome. To test this hypothesis, we used a dataset in which participants share a monetary reward with a close friend (social reward) compared to sharing a monetary reward with a stranger (non-social reward) in a gambling context (dataset 10 (Fareri et al., 2012b)), and found that the trust model was unable to discriminate between shared social rewards from shared non-social rewards (N=20; Table S1; accuracy = 55%, *p* = 0.415; Figure 3B-4) (Fareri et al., 2012b). Thus, contrary to our hypotheses, our results revealed that regardless of reward types, the trust model has no clear association with different aspects of reward processing across all four datasets (Figure 3A-2).

### Specificity of trust model

There are many other potential psychological aspects of the trust experience that can be evaluated using this neurometric approach. First, it is possible that people may be deliberating between keeping the money or investing it in their partner, which may cause decision conflict (Rand et al., 2012; Venkatraman et al., 2009). We tested this hypothesis by applying the model to a stop signal task (dataset 11; N=19; Table S1) (Xue et al., 2008), in which participants are instructed to override a prepotent response, and found that the trust model was unable to discriminate between the successful inhibition and inhibition failure conditions (accuracy = 58%, *p* = 0.322; Figure 3B-5). This suggests that response conflict is not related to the construct of trust, perhaps because cognitive control is common to both decisions to trust and distrust. Decisions to trust may also require social cognition to consider the other player’s mental states such as their beliefs, preferences, and financial outcomes. In order to demonstrate the specificity of the trust construct, we additionally tested our model on several datasets probing distinct aspects of social cognition. We found that the trust model did not generalize to perceptual judgments such as familiarity, in which participants judged whether a face is familiar or unfamiliar to the participants (dataset 12; N=16; accuracy = 63%, *p* = 0.230; Figure 3B-5; Table S1) (Wakeman and Henson, 2015). We also found that the trust model did not generalize to the classification between viewing social and non-social scenes in dataset 13 (N=36; accuracy = 47%, *p* = 0.685; Figure 3B-5; Table S1) (Chen et al., 2019). In addition, we tested if the trust model was similar to self-referential cognition in a task in which participants made self-referential judgments or perceptual judgments (e.g., type of font) to a variety of trait adjectives (dataset 14; N=27; accuracy = 40%, *p* = 0.873; Figure (B-5); Table S1) (Chen et al., 2015, 2013). Lastly, we also found that the trust model did not generalize to discriminate between engaging in theory-of-mind (ToM) reasoning in a task in which participants view shapes engaging in social interactions compared to when the shapes moved randomly (NoToM condition; dataset 15; N=484; accuracy = 52%, *p* = 0.197; Figure (B-5); Table S1) (Barch et al., 2013). Together, these findings indicate that the trust model was not associated with cognitive control, social perception, self-referential processing, or theory of mind processes (Figure 3A-2).

### Trust Nomological Network

The previous analyses demonstrated how our multivariate trust model generalizes to a variety of datasets that may contain overlapping psychological processes to the construct of trust. However, the neurometric approach does not necessarily require training a predictive model for each psychological construct, but can also leverage covariation in spatial patterns. In the next set of analyses, we explored the nomological network of psychological states captured by all 15 datasets we used in the model training and validation procedures by computing the spatial similarity of patterns of brain activity elicited by each of the different experimental tasks. This is akin to correlating how questionnaires relate to other questionnaires in more traditional psychometric analyses. We used three different approaches to perform these analyses.

First, we were interested in characterizing the relationship between all of the data at the subject level. It is possible that subjects may vary considerably within a task, or that different tasks might group together. Because these brain maps contain a high number of features (n∼328k) we used Uniform Manifold Approximation and Projection (UMAP) (McInnes et al., 2018), a nonlinear dimensionality reduction technique, to visualize relationships between each subject from all fifteen datasets (N=1,484) based on the spatial similarity of whole-brain activation patterns. This approach projects the data into a lower two-dimensional space while preserving local and global distance. A qualitative assessment of the projection depicted in Figure 4A indicates that there is quite a bit of individual variability within some tasks like the trust experiments, and more spatial consistency in other tasks such as reward, in which individuals tend to group together. The spatial pattern of participants in the trust condition were closer to those of beliefs of safety, feelings of anticipated no-reward or loss, and feelings of neutral or positive affect. In contrast, whole-brain patterns of participants’ activity in the distrust condition were closer to those of beliefs of risk, feelings of reward, and feelings of negative affect. These findings are largely consistent with our model validation results, with the exception that the distrust data were more similar to reward while the trust data were closer to the no-reward/loss conditions. These discrepancies may be attributed to the non-linear nature of the UMAP algorithm, which emphasizes local distances, which could be more subtle in the higher dimensional space of the brain model. For example, models trained using supervised approaches such as SVM tend to be more spatially diffuse as a consequence of the strong regularization required to estimate parameters of a model that has more features (e.g., voxels) than observations (e.g., images).

**Figure 4.**
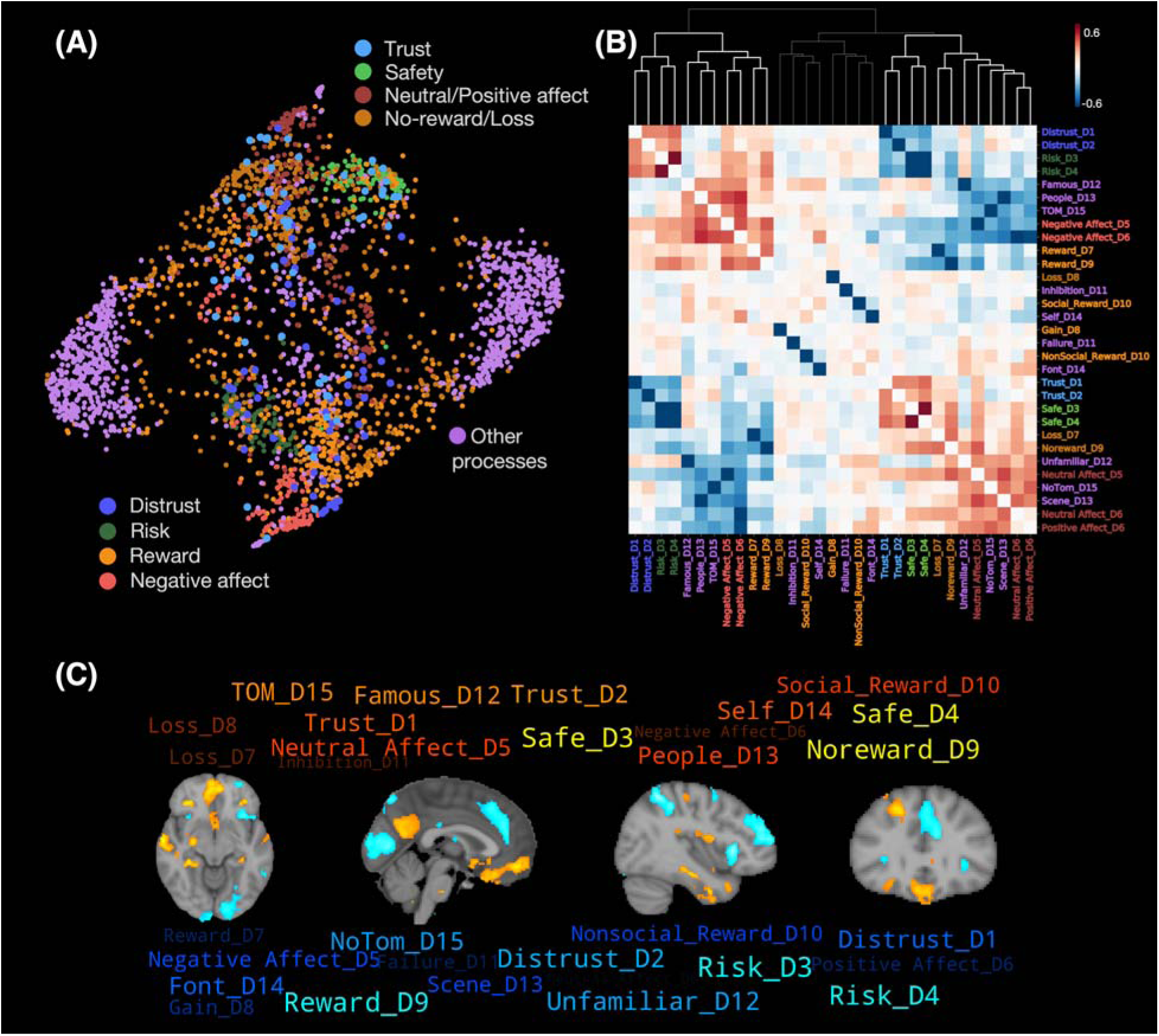
Spatial pattern similarity across all brain data from fifteen datasets. (A) Visualization of spatial similarity of whole brain maps from 1,484 participants from 15 datasets using UMAP nonlinear manifold learning. Each dot represents a beta map from each participant. Parameters were arbitrarily selected to aid in visualization (n-neighbors=50, minimum distance=0.001). Trust was more similar to safety, no-reward, neutral and positive affect; whereas distrust was more similar to risk, reward, and negative affect. (B) Hierarchical clustered heatmap of correlation across the mean spatial pattern from each condition (31 conditions from 15 datasets) also revealed similar findings as above. The dendrogram colors indicate how the tasks cluster together. (C) Independent component analysis (ICA) showed that the two safety, the two trust, three no-reward/loss conditions, and one neutral affect condition loaded positively on the first component, while the two risk, the two distrust, three reward conditions, and two negative affect conditions loaded negatively on this component. See Figure S1 for the other two components

Second, we were interested in examining the relationship between each of the conditions across tasks. We used a more traditional linear approach to quantify the overall spatial similarity of the average whole-brain brain activity patterns within each dataset’s condition. We averaged brain activity across participants for each condition of each task and then computed the pairwise spatial similarity of these maps using Pearson correlations and clustered these maps using hierarchical clustering. These findings are consistent but complementary to the UMAP visualization (Figure 4B). Overall, we found that the datasets group together in at least 3 clusters, with two of these clusters including many tasks that appear to have similar patterns of brain activity. In one of these clusters, the spatial patterns between the two trust conditions were highly similar to each other (r = 0.33), the trust condition in dataset 1 was similar to the two BART safety conditions (r = 0.33 for dataset 3 and r = 0.38 for dataset 4, respectively), and the trust condition in dataset 2 was also similar to the two BART safety conditions (r = 0.23 for dataset 3 and r = 0.32 for dataset 4, respectively). In contrast, in the other cluster, we find that the spatial patterns between the two distrust conditions were highly similar to each other (r = 0.33), the training distrust condition in dataset 1 was similar to the two risk conditions (r = 0.33 for dataset 3 and r = 0.38 for dataset 4, respectively), and the validation distrust condition in dataset 2 was also similar to the two risk conditions (r = 0.23 for dataset 3 and r = 0.32 for dataset 4, respectively). In addition, the safety condition in dataset 4 revealed similar patterns to neutral emotion conditions (r = 0.29 for dataset 5 and r = 0.26 for dataset 6), and the risk condition in dataset 4 was similar to negative emotion conditions (r = 0.29 for dataset 5 and r = 0.25 for dataset 6, respectively; Figure 4B).

Third, we investigated whether there was any evidence of regional covariation across the experimental tasks. All of our analyses throughout the paper have focused on whole-brain activity, but it is possible that specific regions may reflect more specific psychological processes that could be present across many tasks. This analysis is conceptually similar to a factor analysis across different questionnaires. We used independent component analysis (ICA) to reduce the spatial patterns of brain activity across datasets into latent components. Since findings from the hierarchical clustered heatmap showed three prominent clusters, we selected three components for ease of exposition and found that three components explained 51% of the total variance using a principal components analysis (PCA). The first ICA component revealed a spatial pattern similar to our trust model learned by the SVM, r=0.22. Consistent with the results described above, we found that the two trust conditions, the two safety conditions from the two BART tasks, three no-reward/loss conditions, and one neutral affect condition loaded positively on this component, while the two distrust conditions, the two risk conditions from the two BART tasks, three reward conditions, and one negative affect condition loaded negatively on the first component (Figure 4C, Figure S1). In contrast, the other two components were not spatially similar to the trust model (second component, r = 0.09; third component, r = 0.05).

Together these results provide a clear network of relations between the psychological processes elicited by the various tasks that complement the results of the trust model validation analyses. The construct of trust is more closely related to beliefs of safety, anticipated no-reward, and non-negative affect, but not to beliefs of risk, anticipated reward, or negative affect.

## Discussion

In this study, we sought to create a model of trust based on patterns of brain activity elicited during an interpersonal investment task. We employed a neurometric approach (Chang et al., 2015, 2013; Krishnan et al., 2016; Wager et al., 2013; Woo et al., 2017) to characterize this model by assessing its reliability and validity using multiple previously published open datasets. This model leverages reliable patterns of brain activity and is sensitive to detecting a psychological state of trust that generalizes to new subjects, scanners, and variants of the investment game task. In addition, we also assessed the validity of our model (Campbell and Fiske, 1959). Prior work has primarily relied on establishing face validity by demonstrating that regions associated with a construct (e.g., vmPFC) have a reliable independent contribution to the prediction (Chang et al., 2015; Krishnan et al., 2016; Wager et al., 2013). However, directly interpreting the weights of linear models can potentially be misleading as each voxel’s contribution to the prediction is considered in the context of all other voxels. The contributions from voxels that covary may be downweighted when using regularization and voxels may be helping explain a signal that is unrelated to the construct of interest (e.g., denoising) (Kriegeskorte and Douglas, 2019). An alternative approach based on the principles of construct validity attempts to triangulate a construct by establishing its convergent and discriminant validity with related and distinct constructs probed using multiple methods (Campbell and Fiske, 1959; Kragel et al., 2018). This has also been described as establishing a “nomological network” (Cronbach and Meehl, 1955) and identifying the “receptive field” of a model (Woo et al., 2017). We assessed the ability of our trust model to discriminate task conditions across a variety of potentially related psychological constructs elicited using many different types of tasks across 15 previously published datasets, which includes data from over 1,484 participants.

Overall, we found that our brain model of trust was associated with a distinct signature of related psychological processes. First, previous work has established that trust reflects dynamic beliefs about the likelihood of a relationship partner overcoming self-interest and reciprocating (Chang et al., 2010; Delgado et al., 2005). Beliefs about the likelihood of observing either outcome can be modeled as a probability distribution where stronger beliefs of the likelihood of the balloon exploding or not indicate greater certainty while uncertainty arises when either outcome is believed to be likely. Across two separate experiments exploring risky decision-making, our trust model is reliably associated with more safe relative to risky decision contexts. Consistent with these results, we also found that the average pattern of activity when making decisions to trust were spatially similar to average brain activity when making decisions in a safe context, while decisions to not trust were more spatially similar to making decisions in a risky context. These findings indicate that decisions to trust share psychological processes with other decision contexts associated with greater certainty of safety. Second, we find support for the hypothesis that trust requires overcoming concerns of potential betrayal (Bohnet and Zeckhauser, 2004; King-Casas et al., 2008). We find that our trust model is reliably negatively associated with the psychological experience elicited from viewing negative arousing images relative to viewing neutral or positive images. We did not observe a significant relationship with differences between positive vs neutral images indicating that it is neither positive nor neutral images driving this effect. Moreover, pattern similarity analyses revealed that viewing negative images correlated with the risky decision condition, while the neutral images correlated with the safety decisions. These findings are consistent with a betrayal-aversion account. It has been hypothesized that people may choose to keep their money and avoid investing in a relationship partner not just because they don’t want to lose their money, but also because they want to avoid feeling betrayed by another person (Bohnet and Zeckhauser, 2004). Of course, viewing negative arousing images is hardly the same thing as being betrayed and we believe this finding should be further substantiated in future work. Third, contrary to our predictions, we found no evidence that trust is associated with experiencing or anticipating a future reward. We tested our trust model on 4 distinct tasks probing the anticipation and experience of reward and found no indication that trust was related to reward or its anticipation. We think this is particularly important as it has been often assumed that the main motivation for trusting a relationship partner in the trust game is because the expected value is higher (Camerer, 2003; Chang et al., 2010; Fareri et al., 2015). Our findings suggest that it is not the reward, but rather the probability calculus that may be driving decisions to trust. Finally, we also find that trust does not appear to be related to overcoming a prepotent tendency to be selfish, which would recruit cognitive control. Nor does it appear to be involved in social perceptual judgments such as whether an image is a person or an object, if a person has been seen before or is new, or reasoning about another agent’s perspective. We also find no evidence suggesting that trust involves self-referential processing such as considering self-other relative payoffs (Sul et al., 2015). Importantly, we provide additional converging evidence for these analyses using a data-driven approach that uses ICA to identify specific regions that co-vary across all of the datasets included in the paper. One of the components identified using this approach is highly spatially similar to the brain activity pattern identified using our supervised approach with the trust game datasets.

There are several important considerations when interpreting our results. First, we made no assumptions about potential brain regions that may be involved in the psychological experience of trust and chose to utilize a whole-brain approach when training our model (Jolly and Chang, 2021). This demonstrates which regions independently and additively contribute to the prediction. However, it is highly likely that brain activity may be highly collinear, which may lead to instability of the model weights (Haufe et al., 2014; Kriegeskorte and Douglas, 2019), which is addressed by the use of regularization. To mitigate these concerns, we used a bootstrap approach to iteratively retrain the model using different subsets of the data and found that the regions with the largest weights were highly consistent (r=0.91) and also a completely data-driven approach to identified a similar pattern of regional covariation across studies. Future work may consider additionally exploring different types of spatial feature selection (Jolly and Chang, n.d.). Second, our model is currently ignoring interactions between brain regions, which may be an important signature of the trust construct. This might be explored in the future by training new models using functional connectivity or interactions between brain regions. Third, our model is also agnostic to individual differences. We have established that the model generalizes to new participants, but it is not currently able to assess variations in potential motivations (e.g., risk-aversion vs betrayal-aversion). Future work might use multivariate methods for probing individual differences such as intersubject representational similarity analysis (Chen et al., 2020; Finn et al., 2020; van Baar et al., 2019).

In this study, we define trust as the psychological state of assuming mutual risk with a relationship partner to attain an interdependent goal in the face of competing temptations, which is elicited in the context of an interpersonal investment game. Our neurometric results provide evidence indicating that the psychological processes associated with decisions to trust include beliefs that the relationship partner is likely to reciprocate and that decisions to avoid trusting a partner are associated with greater negative affect consistent with the notion of betrayal aversion. Like all brain imaging research, this work relies on the fidelity of eliciting specific psychological processes in experimental paradigms that are conducive to neuroimaging data collection contexts. For example, broad perceptual processes, and higher order cognitive processes such as face perception, language, working memory, and cognitive control appear to be reliably elicited across a variety of paradigms. However, it is considerably more difficult to precisely and reliably elicit more nuanced social and affective processes such as altruism, emotional feelings, and different types of social cognition. Our results are completely dependent on the reliability and validity of the tasks, most of which have never been formally established. We do not believe this had a notable impact on our results, as the majority of the tasks we included are the canonical tasks used to elicit specific processes employed by researchers all over the world and are often included in large scale data collection efforts such as the human connectome project. However, these constructs could potentially be further refined in future work as experimental paradigms that can elicit more subtle nuances of related psychological constructs become publicly available. For example, an interesting question is how the psychological processes measured in a trust game might relate to related constructs such as processes associated with quick perceptual judgments of facial trustworthiness (van ’t Wout and Sanfey, 2008). We see construct validity as an iterative process that cannot be fully addressed by a single paper, but instead will require continued refinement as more data sets become available in the future (Woo et al., 2017). Our ability to test additional hypotheses were impeded by numerous issues we encountered in neuroimaging data sharing. For example, data from previously published datasets are not always accessible due to issues with data backup procedures, meta-data and data formats, or lack of responses to data sharing requests to authors. We hope that changing norms for data sharing advanced by open science initiatives, will result in the broader research community adopting better data preservation practices to facilitate cross-study construct validation efforts.

In summary, using 15 datasets, we establish a neurometric-based construct validity of trust. This model is stored as a three-dimensional brain image that contains a recipe for how to linearly combine information from each voxel in the brain(Jolly and Chang, 2019). Importantly, this model generalizes beyond the specific subjects, scanner, or experimental paradigm and can easily be shared with other researchers (Woo et al., 2017). In addition, we move beyond a reverse inference approach (Poldrack, 2006) in interpreting the psychological processes associated with trust based on which regions contribute to the prediction (Chang et al., 2015, 2013), to a more quantitative construct validity approach. These analyses support several previous accounts of trust, but importantly rule out a reward-based motivation. This provides a proof of concept that brain activity can be used to make inferences about a psychological process beyond self-report or behavioral observations. We believe this general approach could be applied to any other psychological constructs that can be measured using patterns of brain activity.

## Methods

### fMRI Dataset

#### Trust model training datasets

The training datasets (dataset 1-1 and 1-2) for the trust model contained data from two published studies (Fareri et al., 2015, 2012a). In dataset 1-1, 17 participants played an iterated trust game with three different trustees while undergoing fMRI in a 3T Siemens Allegra scanner (TR=2000ms; TE=25ms)(Fareri et al., 2012a). Participants were endowed with one dollar and on each trial decided whether to invest this money in the other trustee (i.e., trust) or keep it (i.e., distrust). Decisions to trust resulted in the one dollar investment being multiplied by a factor of three. The trustee then decided whether to keep all three dollars, or share half of the return on the investment back to the participant (i.e., $1.50). In dataset 1-2, 23 participants also played a similar iterated trust game with two different trustees while undergoing fMRI in a 3T Siemens Magnetom Trio scanner (TR=2000ms; TE=30ms) (Fareri et al., 2015).

In total, there were 40 participants from dataset 1-1 and 1-2 in the current study. We focused our analysis only on the decision epoch when participants made decisions to either trust or distrust. fMRI data were analyzed using a combination of custom scripts (https://github.com/rordenlab/spmScripts) for SPM12 and FSL (v5.09; FMRIB). We performed standard preprocessing in SPM (motion correction, brain extraction and coregistration, slice time correction). Motion artifact was removed using ICA-AROMA in FSL (Pruim et al., 2015). Functional data were smoothed using a 5mm kernel in FSL. Each condition was modeled as a separate regressor in a general linear model (GLM). This included a regressor modeling each of the decision types (trust or distrust) and the different possible decision outcomes (though these data were not the focus of the present manuscript). The GLM resulted in a trust whole-brain beta map and a distrust whole-brain beta map for each trustee (detailed preprocessing and GLM steps see (Fareri et al., 2015, 2012a)). We then averaged the beta maps across partner types within each participant to generate a trust and distrust beta map. The trust and distrust beta maps were then mean-centered by subtracting the mean across all voxels within each map(Misaki et al., 2010) and used to train the trust model.

#### Trust model validation dataset

The validation dataset (dataset 2) contained data from 17 participants (mean age = 20.6 years, SD=1.49; 24% female) who participated in a repeated trust game while undergoing fMRI in a 3T Philips Achieva scanner (TR = 2200 ms, TE = 30 ms, FOV = 220 × 220 × 114.7 mm; see (Schreuders et al., 2018) for more details about the sample and scanning parameters). All participants provided informed consent and the study was approved by the institutional review board at Leiden University Medical Center. Participants were instructed to play a trust game with three different targets, including a friend, an antagonist, and an anonymous peer. The game was designed to be slightly more similar to a prisoner’s dilemma in that both players made their decisions simultaneously. Unlike a traditional trust game, participants received information about their partner’s decisions regardless if they chose to share or keep. However, the responses from these targets were pre-determined by the computer and not the actual partner. Similar to dataset 1, we also focused our analysis on the decision epoch when participants made either a trust or distrust decision. Image pre-processing and analysis was conducted using SPM8 software (www.fil.ion.ucl.ac.uk/spm) implemented in MATLAB R2010 (MathWorks). Pre-processing included slice-time correction, realignment, spatial normalization to EPI templates, and smoothing with a Gaussian filter of 8 mm full-width at half maximum (FWHM). The fMRI time series were modeled by a series of events convolved with a canonical hemodynamic response function (HRF). The data was modeled at choice and feedback onset as null duration events. During decision-making the choice events (i.e., trust and keep decisions) were modeled for each of the three partner types. These modeled events were used as regressors in a general linear model (GLM) with a high pass filter using a discrete cosine basis set with a cutoff of 120 seconds. The GLM resulted in a trust whole-brain beta map and a distrust whole-brain beta map for each target. We then computed the mean trust whole-brain beta map across all three targets and repeated the same procedure for computing the mean distrust whole-brain beta map within each participant. We then mean-centered values across all voxels within each of the beta maps for all participants, and the mean-centered beta maps were used as a novel trust dataset for brain model validation.

#### Safety datasets

In order to test whether trust is associated with beliefs of safety, we used the Balloon Analog Risk Task (BART), in which one condition probes beliefs of risk and another probes beliefs of safety in this study. The BART aims to elicit naturalistic risk-taking behaviors, and each participant received two conditions in the fMRI scanner. In the risk condition, each inflation of balloons is a risky choice (pump), whereas inflating balloons in the safe condition is not a risky choice (control pump). In dataset 3 (OpenfMRI ds000001) (Schonberg et al., 2012), there are 15 healthy participants who underwent the two conditions in a 3T Siemens Allegra MRI scanner. FMRIPREP(Esteban et al., 2019) was used for brain data preprocessing, and the steps included motion correction, skullstripping and coregistration to T1 weighted volume, applying brain masks, realignment, and normalization. Data were spatially smoothed with a 6mm FWHM Gaussian kernel and first level analyses were performed using FEAT in FSL (v 6.0.4). For trials in the risky condition, the risky inflation and the other two task-related regressors were modeled separately in the GLM. For trials in the safe condition, the safe inflation and the other two task-related regressors were also modeled separately in the GLM. For each participant, the GLMs resulted in a risk inflation whole-brain beta map and a safe inflation whole-brain beta map. We then mean-centered values across all voxels within each beta map for all participants, and these mean-centered beta maps were used as data representing the risk condition and safety condition in the generalization testing.

In dataset 4 (OpenfMRI ds000030) (Poldrack et al., 2016), there are 124 healthy participants who also underwent the two conditions in a 3T Siemens Allegra MRI scanner. This dataset was collected by the Consortium for Neuropsychiatric Phenomics as part of a larger study focused on understanding the dimensional structure of cognitive processing in healthy individuals and those diagnosed with neuropsychiatric disorders (Poldrack et al., 2016). All participants gave written informed consent following procedures approved by the Institutional Review Boards at UCLA and the Los Angeles County Department of Mental Health. Imaging data were collected on a Siemens Trio 3T scanner using EPI sequence (TR=2000ms, TE=30ms, flip angle=90°, matrix=64 x 64, FOV=192mm, 34 oblique slices). Data was accessed from OpenNeuro (https://openneuro.org/datasets/ds000030/versions/1.0.0) and preprocessed using fMRIPrep (Esteban et al., 2019), followed by a spatial smoothing with a 6mm FWHM Gaussian kernel. A risk inflation (accept pump) and a safe inflation (control pump), along with the other 24 motion parameters (6 realignment parameters, their squares, derivatives and squared derivatives), motion splikes, global signal-intensity spikes more than 3 SDs above the mean of the intensity between TRs were modeled in the GLM by using nltools (Chang et al., 2018). A risk inflation (accept pump) and a safe inflation (control pump), along with the other seven regressors were modeled in the GLM. For each participant, the GLM resulted in a risk inflation whole-brain beta map and a safe inflation whole-brain beta map, and we then mean-centered values across all voxels within each beta map for all participants. These mean-centered risk inflation beta maps and safe inflation beta maps were then used as data representing the risk condition and safety condition in the generalization testing.

#### Affect datasets

Two affect datasets were included in the current study. Dataset 5 came from the PINES dataset (Chang et al., 2015), which was an open dataset on Neurovault (https://identifiers.org/neurovault.collection:503). In this dataset, participants were asked to view numerous negative and neutral-valenced pictures from the international affective picture system (IAPS), and then rated how negative they felt from 1 (neutral) to 5 (most negative). Details of the experimental design were described in previous studies (Chang et al., 2015; Gianaros et al., 2014). Among these participants, this current study only used data from those (N = 93) whose ratings had 1 (neutral) and 5 (most negative). The fMRI data was collected in a Siemens Trio 3T scanner (TR= 2000 ms, TE=29ms), and then preprocessed by SPM8, including unwarping, realignment, coregistration, normalization, spatial smoothing with a 6 mm FWHM Gaussian kernel and high pass filtering (180 sec cutoff). Then five separate regressors indicating different rating levels (1 to 5) were modeled in the GLM for each participant as well as 24 covariate regressors modeled movement effects (6 realignment parameters demeaned, their 1st derivatives, and the squares of these 12 regressors). Since our goal was to compare the neutral and negative condition, only the neutral (rating = 1) beta map and the negative (rating = 5) beta map were included in the current study. We then mean-centered values across all voxels within each of the above two kinds of beta maps for all participants. These mean-centered neutral and negative beta maps were taken as data in the generalization testing.

In Dataset 6, fifty-six participants were recruited to complete an emotional scene task (Hsiao et al., 2023). In this task, participants were asked to make indoor/outdoor categorization judgments on scenes in a block design. Each block lasted 15 seconds and consisted of six emotional scenes with the same emotional valence. Each emotional-scene block alternated with a 15-sec fixation block, and each participant went through five blocks for each of three different valences, including positive, neutral, and negative valence. The emotional valence of the scenes used in each condition were selected from the IAPS and have been validated in a previous fMRI study (Wagner and Heatherton, 2012). The fMRI data was collected in a Philips Intera Achieva 3T scanner (TR = 2500 ms, TE = 35 ms), and then preprocessed by SPM8, including slice timing correction, unwarping, realignment, coregistration, normalization, and spatial smoothing with a 6 mm FWHM Gaussian kernel. The positive, neutral, and negative valence conditions were then modeled separately in the GLM for each participant. The GLM resulted in a positive, neutral, and negative emotion beta map from each participant, and we then mean-centered values across all voxels within each beta map for all participants. These mean-centered beta maps were used in the current study, representing three different emotional-valence conditions in the generalization testing.

#### Reward datasets

Four reward anticipation fMRI datasets were used in the current study. Dataset 7 comes from the Human Connectome Project (Barch et al., 2013), and the reward anticipation task used in this dataset is the Card Gambling task (Delgado et al., 2000). In this task, participants were asked to guess whether the number on a mystery card is greater or smaller than five. Participants would receive a reward of one dollar if the number is greater than five; by contrast, they would lose fifty cents if the number is smaller than five. In total, fMRI data from 490 participants were collected and preprocessed with the HCP fMRIVolume pipeline (Glasser et al., 2013). The preprocessing steps included gradient unwarping, motion correction, fieldmap-based EPI distortion correction, coregistration, normalization, and spatial smoothing with a 4 mm FWHM Gaussian kernel. The reward and loss conditions were then modeled in the GLM. The GLM resulted in a reward beta map and a loss beta map within each participant, and we then mean-centered values across all voxels within each beta map for all participants. These mean-centered reward and loss beta maps were then used as data representing the reward condition and non-reward/loss condition in the generalization testing.

In dataset 8, eighteen participants completed a reward/loss anticipation task while undergoing scanning in a 3T Siemens Trio scanner (Zhang et al., 2017). In this task, different cues were shown on the screen indicating different amounts of monetary reward or loss. After the cue phase, an outcome phase occurred, indicating the actual amount of reward and loss. The monetary reward or loss amounts were equally sampled from [1, 5, 20, 100]. The data were preprocessed using BrainVoyager QX 2.8 and NeuroElf V1.1, including: motion correction, slice timing correction, high-pass filtering, normalization, and spatial smoothing with a 6 mm FWHM Gaussian kernel. The cue and outcome phases with different levels were modeled separately in the GLM for each participant. Only the maximal-reward (i.e., a gain of $100) and maximal-loss (i.e., a loss of $100) beta maps from each participant were used in the current study, and we then mean-centered values across all voxels within each beta map for all participants. These mean-centered beta maps would represent the reward condition and non-reward/loss condition in the generalization testing.

In dataset 9 (OpenNeuro ds003242) (Tomova et al., 2020), twenty-nine participants underwent a monetary incentive delay task (Knutson et al., 2000; Krebs et al., 2011) in an fMRI scanner. Before the task, participants were asked to memorize five abstract art images, and these familiar images were then taken as cues in the reward condition. In the reward condition, after a familiar image was shown on the screen as a reward cue, a number ranging from 1 to 9 was shown and participants had to respond whether the number was larger or smaller than 5. If participants responded fast enough (< 500 ms), they would receive a reward of one dollar. In the other condition, the non-reward condition, after a new abstract art image was shown as a non-reward cue, a number was also shown on the screen and participants were also asked to respond whether the number is greater or smaller than 5. However, the responding performance in the non-reward condition was not associated with any reward. FMRIPREP (Esteban et al., 2019) was used for brain data preprocessing, and the steps included motion correction, skullstripping and coregistration to T1 weighted volume, applying brain masks, realignment, and normalization. Data were spatially smoothed with a 6mm FWHM Gaussian kernel and first level analyses were performed using FEAT in FSL (v 6.0.4). We modeled the reward condition and non-reward conditions as separate regressors in a univariate GLM, along with 24 covariate regressors modeling movement effects (6 realignment parameters demeaned, their 1st derivatives, and the squares of these 12 regressors), a 128 sec high pass filter using a discrete cosine transform, and separate scanner spikes based on frame differences that exceeded 3 standard deviations. For each participant, the GLM resulted in a reward beta map and a non-reward beta map, which were then mean-centered across all voxels within each beta map for all participants. These mean-centered reward and non-reward beta maps were then used as data representing the reward condition and non-reward/loss condition in the generalization testing.

In dataset 10, twenty participants underwent a shared social reward task in a Siemens 3T Allegra scanner (Fareri et al., 2012b). In this task, each participant played a card guessing game for shared monetary rewards with three different partners, including a close friend, a stranger (confederate) they met at the scan session, and a computer. Each participant was scanned in the scanner and took turns in playing or observing their partner (friend or stranger) who were in the scanner control room across scanning blocks. They needed to guess whether the value of a playing card was greater or smaller than 5. They received a shared monetary gain of $4 with their partner for each correct trial, but received a shared monetary loss of $2 for each incorrect trial. In this study, we defined the shared monetary reward condition with a friend as the social reward condition and the shared monetary reward condition with a stranger as the non-social reward condition. FMRIPREP(Esteban et al., 2019) was used for brain data preprocessing, and the steps included motion correction, skullstripping and coregistration to T1 weighted volume, applying brain masks, realignment, and normalization. Data were spatially smoothed with a 6mm FWHM Gaussian kernel and first level analyses were performed using FEAT in FSL (v 6.0.4). A GLM was computed including regressors modeling the choice (3 regressors) and outcome phases (6 regressors) of the card game for each partner. Regressors of no interest were included modeling the choice and outcome phases of missed trials. Confound regressors modeling trial-by-trial framewise displacement, motion in six planes (translation, rotation), the first six principal components derived from aCompCor capturing physiological noise and cosine basis functions were included for each participant in each run. For each participant, the GLM resulted in a shared social reward beta map and a shared non-social reward beta map, which were then mean-centered across all voxels within each beta map for all participants. These mean-centered social reward and non-social reward beta maps were then used as data representing the social reward condition and non-social reward condition in the generalization testing.

#### Other processing datasets

In order to demonstrate the specificity of our trust model, we validated our model on four additional datasets, including cognitive control, familiarity, social cognition and self-referential cognition. To test the domain of cognitive control, in Dataset 11, nineteen participants performed a stop-signal task (SST) in a 3T Siemens Allegra MRI scanner (TR=2000ms, TE=30ms) (Xue et al., 2008). This open dataset is available on both OpenNeuro (ds000007) and Neurovault (https://neurovault.org/collections/1807/). We used data from Neurovault task001, which was a manual SST. For the go trials in this task, participants were asked to press on the right or left button according to whether the letter “T” or “D” was shown on the screen. For stop trials, an auditory tone cue signaling stop was played after the letter being shown with some delay (stop-signal delay; SSD), and participants were asked to inhibit their approaching responses toward the button. Throughout the task, the length of SSD changed according to whether participants succeeded or failed to inhibit their responses in order to maintain the accuracy rate at 50%. Thus, the number of inhibition-success and inhibition-failure trials would be the same, and we would use data from both of these two conditions for further analysis. The data was preprocessed by FSL version 3.3, and the preprocessing steps included coregistration, realignment, motion correction, denoising using MELODIC, normalization, spatial smoothing with a 5 mm FWHM Gaussian kernel, and high-pass filtering. Details about the preprocessing steps were described in the original study (Xue et al., 2008). Four conditions, including go, inhibition-success, inhibition-failure, and nuisance events, were modeled separately in the GLM for each participant. The GLM resulted in a go, inhibition-success, inhibition-failure, and nuisance event beta map, and we then mean-centered values across all voxels within each beta map for all participants. Only the mean-centered inhibition-success and inhibition-failure beta maps were used in the generalization testing.

For the domain of familiarity, in Dataset 12, sixteen participants completed a face-viewing task in a Siemens 3T TRIO scanner (TR=2000ms, TE=30ms) (Wakeman and Henson, 2015). This open dataset is available on both OpenNeuro (ds000117) and Neurovault (https://neurovault.org/collections/1811/), and we used data downloaded from Neurovault. In this face-viewing task, participants were asked to view three different types of faces, including famous, unfamiliar, and scrambled faces. Each trial began with a fixation cross on the screen, and then one of the three types of faces were shown on the screen. Participants were asked to pay attention to all trials throughout the whole experiment. The fMRI data was preprocessed by SPM8, which included slice timing correction, realignment, coregistration, normalization, and spatial smoothing with a 8 mm FWHM Gaussian kernel. Three conditions, including famous, non-familiar, and scrambled faces were modeled separately in the GLM for each participant. The GLM resulted in a famous, non-familiar, and scrambled beta map, and we then mean-centered values across all voxels within each beta map for all participants. Only the mean-centered famous and non-familiar beta maps were used in the current study for the generalization testing.

For the domain of social cognition, in Dataset 13, thirty-six participants completed a scene judgment task in a Philips Intera Achieva 3T scanner (TR = 2500 ms, TE = 35 ms) (Chen et al., 2019). In this task, each participant was asked to make indoor/outdoor categorization judgements on 270 different scenes, including 90 social scenes, 90 non-social scenes, and another 90 food scenes. These pictures have been used in several studies (Powers et al., 2011; Wagner et al., 2013, 2012), and compared to non-social scenes, social scenes have been found to reliably activate brain regions, such as the dmPFC, PCC, and vmPFC (Powers et al., 2011). In each trial, a scene image was shown on the screen for 2000 ms, followed by a 500 ms fixation, and a jitter (range: 0-5000 ms) was followed between each trial. The fMRI data were preprocessed by SPM8, which included slice timing correction, unwarping, realignment, motion correction, normalization, spatial smoothing with a 6 mm FWHM Gaussian kernel. The social, non-social, and food conditions were then modeled separately in the GLM for each participant. The GLM resulted in a social, non-social, and food beta map, and we then mean-centered values across all voxels within each beta map for all participants. Only the mean-centered social and non-social beta maps from each participant were then used in the current study, representing the social and non-social condition for the generalization testing.

For the domain of self-referential cognition, in Dataset 14, twenty-seven participants completed a trait-judgment task in a Philips Intera Achieva 3T scanner (TR = 2500 ms, TE = 35 ms) (Chen et al., 2015, 2013). In this trait-judgment task, participants were asked to make three different targets of judgments, including self-judgment (i.e. does this adjective describe you?), mother-judgment (i.e. does this adjective describe your mother?) and font-judgment (i.e. is this adjective printed in bold-faced letters?) in two different languages. For each trial, a trait adjective word (e.g. smart) was paired with a target word (i.e. SELF, MOTHER, or FONT) and were shown on the screen for 2500ms. Although each trait word was presented once in Mandarin and once in English, the current study only used the three conditions in Mandarin. The fMRI data was preprocessed by SPM8, which included slice timing correction, unwarping, realignment, motion correction, normalization, spatial smoothing with a 6 mm FWHM Gaussian kernel. The self-judgment, mother-judgment, and font-judgment conditions were then modeled separately in the GLM for each participant. The GLM resulted in a self-judgment, mother-judgment, and font-judgment beta map, and lastly we mean-centered values across all voxels within each beta map for all participants. Only the mean-centered self-judgment and font-judgment beta maps from each participant were then used for the generalization testing in the current study.

For the domain of theory of mind, in Dataset 15, 484 participants watched a short 20-second engaging video clip in which objects (circles, squares, and triangles) show interactive behaviors (ToM condition), and they also watched another video clip in which the same objects move randomly (NoToM condition). In total, fMRI data from 484 participants were collected and preprocessed with the HCP fMRIVolume pipeline (Glasser et al., 2013). The preprocessing steps included gradient unwarping, motion correction, fieldmap-based EPI distortion correction, coregistration, normalization, and spatial smoothing with a 4 mm FWHM Gaussian kernel. The ToM and NoToM conditions were modeled as a block design regressor in the GLM for each participant. The GLM resulted in a ToM and a NoToM beta map, and we then mean-centered values across all voxels within each beta map for all participants. Only the mean-centered ToM and NoToM beta maps from each participant were then used for the generalization testing.

### Training and validating a trust model

#### Training model and cross-validation within the training dataset

We used a three stage approach to train our whole-brain multivariate classification model using a linear Support Vector Machine (SVM) by using the predict package from Nltools (Chang et al., 2018). First, we were interested in evaluating how well the model might generalize to new data using a leave-one-subject-out (LOSO) cross-validation procedure, ensuring that every subject served as both training and testing data (Chang et al., 2015). This allowed us to evaluate how a model trained on 39 participants could classify trust or distrust decisions from the left-out participant and provided an estimate of the expected generalizability of the model to similar datasets. Second, we were interested in assessing which voxels most consistently contributed to the trust classification. We used a parametric bootstrap procedure, which involved retraining the model 5,000 times after randomly sampling participants with replacement. The resulting distribution was then converted into a z-value at each voxel, which allowed the map to be thresholded based on a corresponding p-value. We used *p* < 0.005 as the threshold to visualize the most reliable weights to help assess the face validity of the model (Figure 2A). It is important to note that we did not use this thresholded map to perform any inferences. We further computed the spatial intersubject correlation across the models trained on each bootstrapped sample to estimate the approximate consistency of the spatial pattern of weights. This metric can be interpreted similarly to a reliability coefficient, but will be somewhat inflated compared to using completely independent data. Third, we trained the final model using the data from all participants, which is what we ultimately used to test on all other datasets. This model will have the best fitting weights as it was trained on all available data. For all tests, we used a forced-choice accuracy procedure to evaluate the performance of the model. Forced-choice accuracy examines the relative expressions of the model between the two brain images collected from the same participant and is well suited for fMRI as the input images are unlikely to be on the same scale across individuals or scanners (Chang et al., 2015; Wager et al., 2013). We performed hypothesis tests using permutations in which the labels for each image across participants were randomly flipped 10,000 times to generate a null distribution. We were only interested in whether the target condition was significantly greater than the reference condition, so we reported one-tailed tests. We also computed receiver operator character (ROC) curves using forced choice accuracy. An interesting property of forced choice accuracy is that it is equivalent to sensitivity, specificity, and area under the curve (AUC) of the ROC curve.

#### Model validation using an independent testing dataset

In order to examine the validity of the trust brain model in an even more rigorous and unbiased way beyond cross-validation, we evaluated its generalizability to a new test dataset (Dataset 2). This dataset was collected in a different country (The Netherlands), using a different scanner and variant of the trust game. We computed forced-choice accuracy on this dataset based on the spatial similarity of the trust model and each participant’s trust and distrust beta images estimated using a first level GLM, and also calculated an ROC curve to quantify the tradeoff of sensitivity and specificity at different thresholds (Figure 2B).

### Construct validity and specificity of trust model

To evaluate the convergent and discriminant validity of the trust model to other psychological constructs, we tested our trust classification model on other datasets probing distinct psychological constructs, including: beliefs of safety (Dataset 3 and 4), negative affect (Dataset 5 and 6), feelings of anticipated reward (Dataset 7, 8, 9, and 10), cognitive control (Dataset 11), social cognition (Dataset 12, 13, and 15), and self-referential cognition (Dataset 14).

For each dataset, we computed the spatial similarity between the trust multivariate brain pattern and each participant’s beta maps representing the test and control conditions from each task. For example, we evaluated how well the trust model could discriminate between safety and risk in datasets 3 and 4, neutral and negative emotional experience in dataset 5 and 6, positive and negative emotional experiences in dataset 6, anticipated reward and loss in dataset 7, anticipated money gain and loss in dataset 8, anticipated reward and no-reward in dataset 9, social reward and loss in dataset 10, success and failure in cognitive control in dataset 11, familiarity and unfamiliarity in dataset 12, social and non-social viewing in dataset 13, self and non-self referential cognition in dataset 14 as well as ToM and NoToM in dataset 15. We followed the same forced-choice testing procedure outlined above. Assessing the generalizability of the trust model across different datasets in this manner allowed us to demonstrate convergent and discriminant validity of the trust brain model with other psychological constructs.

### Trust Nomological network

In order to assess the trust nomological network, we conducted three different analyses to assess the overall relationships between the 15 datasets based on spatial similarity of the brain imaging data.

First, to qualitatively visualize the similarity of all of the participants (N=1,000) from all 15 datasets, we used Uniform Manifold Approximation and Projection (UMAP), a nonlinear dimensionality reduction technique (Figure 4A; https://github.com/lmcinnes/umap). UMAP attempts to project high dimensional data into a low dimensional space preserving both local and global distance in the feature space using manifold learning (McInnes et al., 2018). It is well suited for visualizing large datasets with high-dimensionality. We first removed subject-specific mean activity across the two condition brain maps within each dataset. We used arbitrarily selected values for the hyperparameters (number of neighbors = 50, minimal distance = 0.001). We note that UMAP is a stochastic technique that is sensitive to the parameters that are selected and will give a slightly different projection depending on the random seed initialization, though the relative distances will be preserved.

Second, to quantitatively assess the overall similarity between the datasets, we averaged beta maps across participants for each condition and computed the spatial similarity across 31 conditions from all datasets using pairwise Pearson correlations. We visualize these relationships using a hierarchical clustered heatmap (Figure 4B).

Third, to determine which regions were common across datasets, we performed an independent component analysis (ICA). This is conceptually similar to performing a factor analysis across questionnaires to identify how each question loads onto latent factors measured by the questionnaires. We created an observation-by-features matrix by vectorizing the whole-brain voxel beta map associated with each condition after averaging across participants, which effectively treats a voxel as a question and a condition as one data observation. We performed a spatial decomposition using ICA across all conditions using the Fast ICA algorithm from Scikit-learn (Pedregosa et al., 2011). We chose to limit our analysis to only identifying 3 components based on the results of our hierarchical clustering analysis (Figure 4B). Three components explained 51% of the total variance as determined by PCA and resulted in distinct spatial patterns. We plot the spatial pattern and loading weights for each condition as a wordcloud in Figure 4C. Full results of the spatial pattern and loading weights for each condition are reported in the supplementary materials (Figure S1).

## Acknowledgements

The authors thank Zhihao Zhang and Ifat Levy for sharing their data (dataset 8) and Wasita Mahaphanit for her comments on the paper. This research was supported by funding from the Chiang Ching-Kuo Foundation for International Scholarly Exchange (no. GS040-A-16 to P.-H.C.), the Young Scholar Fellowship Program by the Ministry of Science and Technology and National Scoence and Technology Council (MOST 109-2636-H-002-006, 110-2636-H-002-004, 111-2628-H-002-004, and NSTC 111-2423-H-002-008-MY4 to P.-H.C.) in Taiwan, National Institute of Health grants (nos. R01MH116026 and R56MH080716 to L.J.C., R15MH122927 to D.S.F.), the McKnight Foundation (to M.R.D.), the Netherlands Organization for Scientific Research (NWO; VENI Grant Number 451-10-021 to B.G.) and the National Science Foundation (NSF CAREER 1848370 to L.J.C.). The code used to perform the analysis in the paper will be available on github pending publication of this manuscript. The trust brain model as well as brain beta images in dataset 1, 2, 6, 10, 13 and 14 will be uploaded to Neurovault pending publication of this manuscript. Datasets 3, 4, 9, 11 and 12 are downloaded from OpenfMRI platform, dataset 5 is downloaded from Neurovault platform, and datasets 7 and 15 are downloaded from Human Connectome Project website.

## Supplementary materials

**Table S1.**
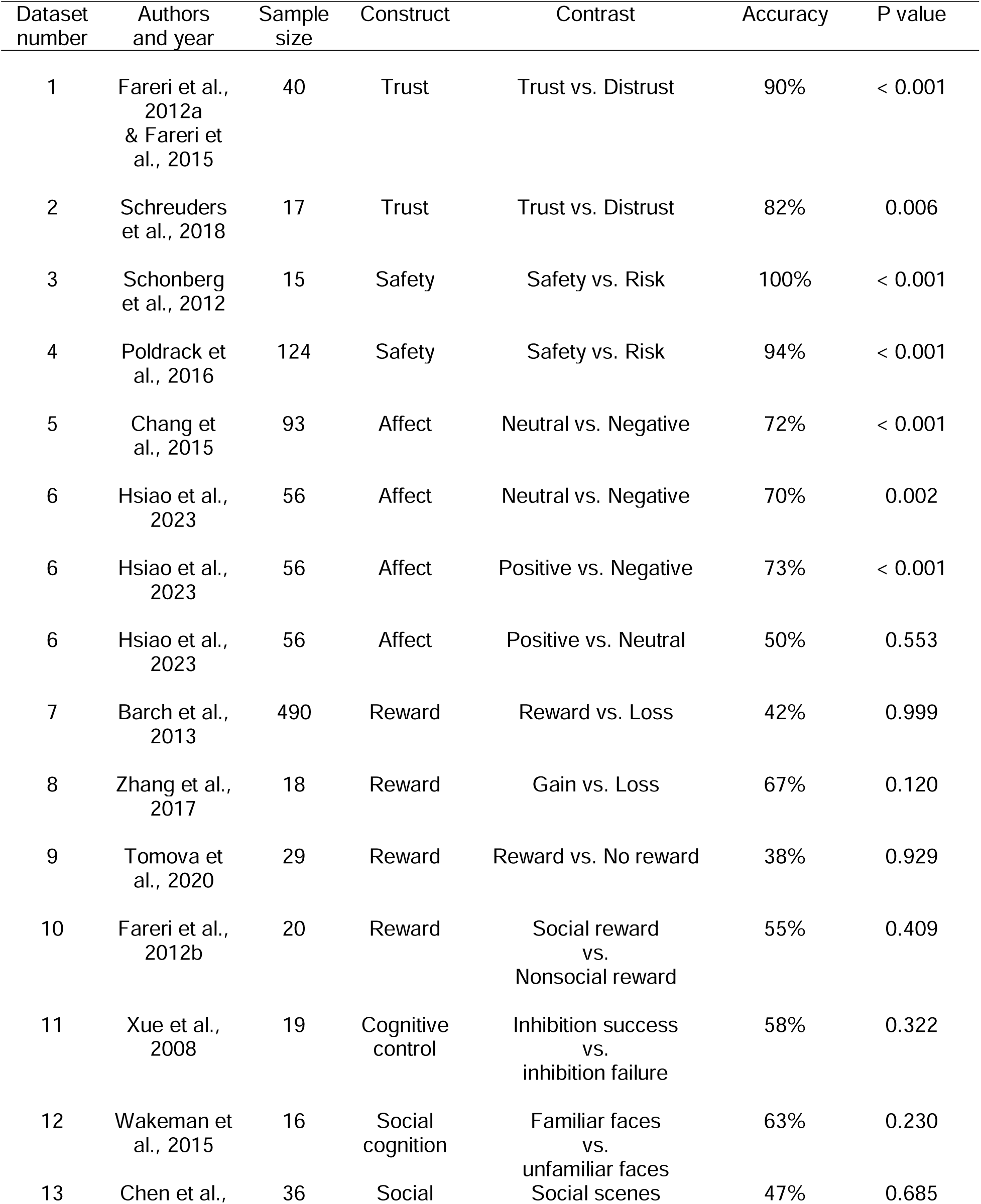
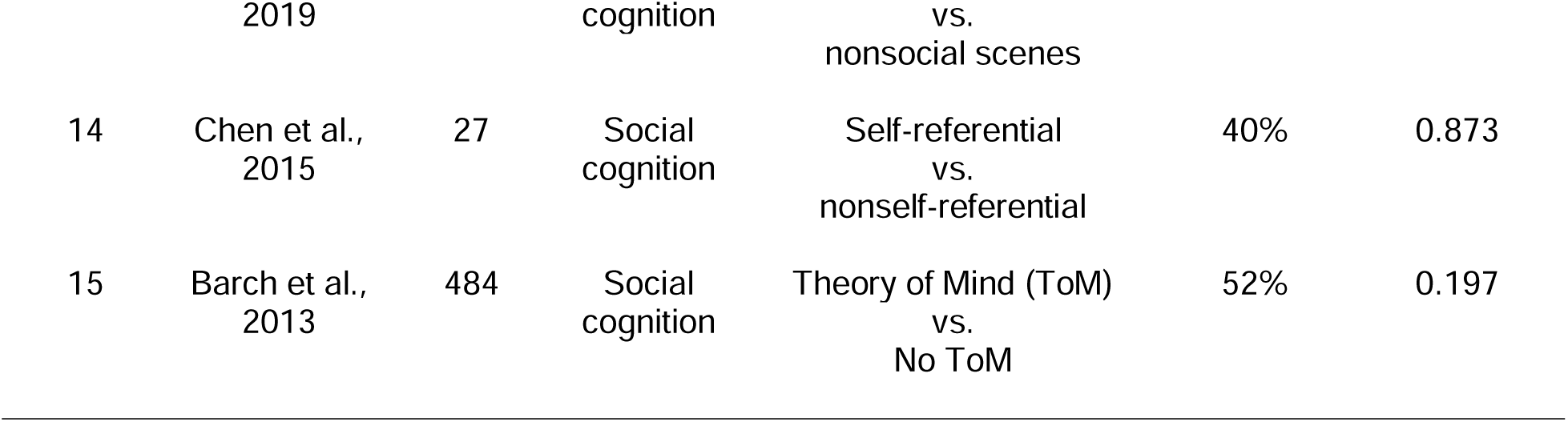
Basic information of each dataset as well as forced-choice classification accuracy and p values for each generalization testing.

**Figure S1.**
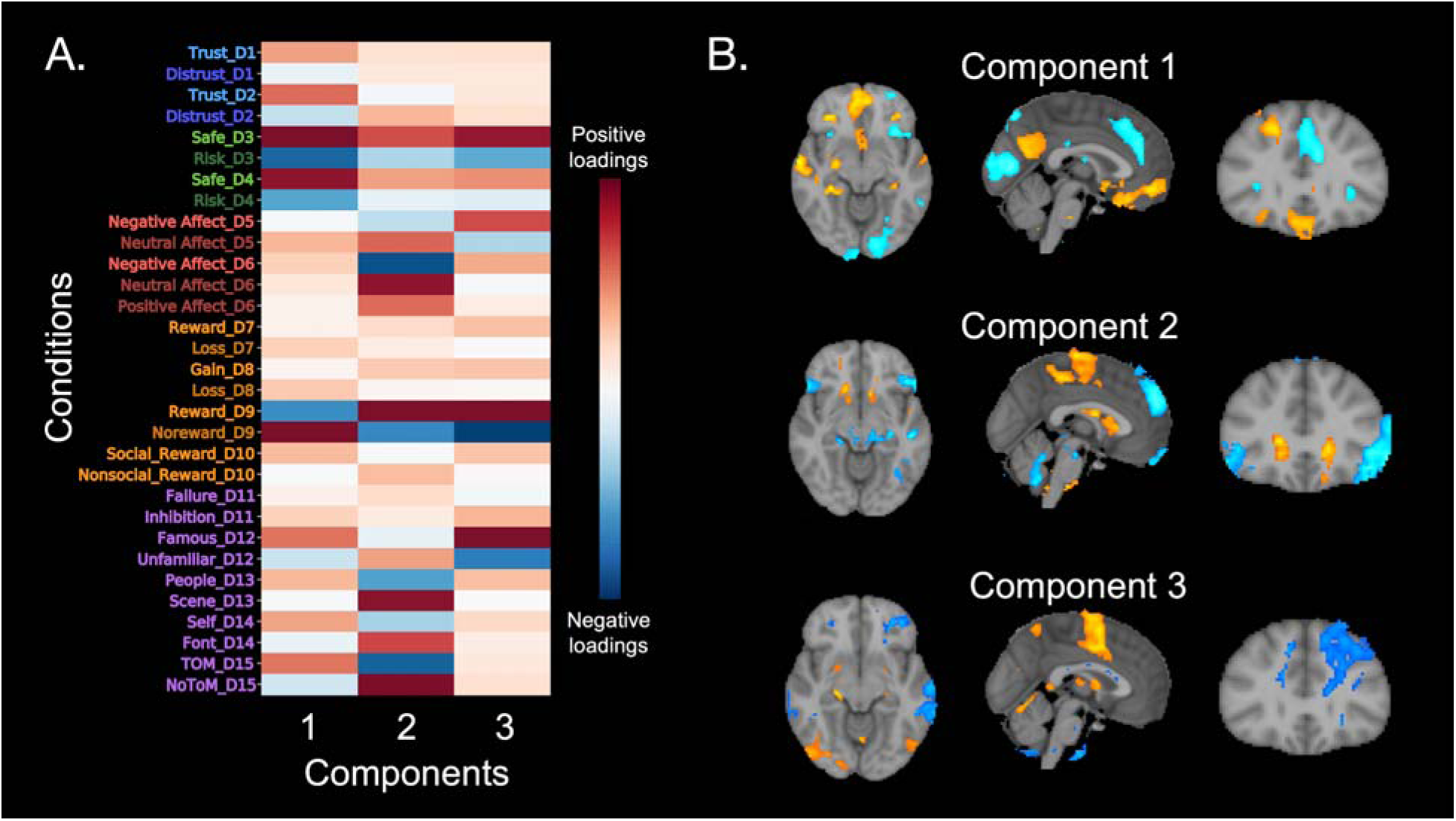
Condition loadings on each of the three ICA components. (A) The two trust conditions, safe conditions, no-reward/loss conditions, and one neutral affect condition loaded positively on the first component. By contrast, the two distrust conditions, risk conditions, reward conditions, and one negative affect condition loaded negatively on the first component. (B) Spatial patterns for each of the three components.

