## Supplementary_figure_table for "Towards a Neurometric-based Construct Validity of Trust"

### Supplementary materials

**Table S1.** Basic information of each dataset as well as forced-choice classification accuracy and p values for each generalization testing.

| Dataset number | Authors  and year | Sample size | Construct | Contrast | Accuracy | P value |
| --- | --- | --- | --- | --- | --- | --- |
| 1 | Fareri et al., 2012a  & Fareri et al., 2015 | 40 | Trust | Trust vs. Distrust | 90% | < 0.001 |
| 2 | Schreuders et al., 2018 | 17 | Trust | Trust vs. Distrust | 82% | 0.006 |
| 3 | Schonberg et al., 2012 | 15 | Safety | Safety vs. Risk | 100% | < 0.001 |
| 4 | Poldrack et al., 2016 | 124 | Safety | Safety vs. Risk | 94% | < 0.001 |
| 5 | Chang et al., 2015 | 93 | Affect | Neutral vs. Negative | 72% | < 0.001 |
| 6 | Hsiao et al., 2023 | 56 | Affect | Neutral vs. Negative | 70% | 0.002 |
| 6 | Hsiao et al., 2023 | 56 | Affect | Positive vs. Negative | 73% | < 0.001 |
| 6 | Hsiao et al., 2023 | 56 | Affect | Positive vs. Neutral | 50% | 0.553 |
| 7 | Barch et al., 2013 | 490 | Reward | Reward vs. Loss | 42% | 0.999 |
| 8 | Zhang et al., 2017 | 18 | Reward | Gain vs. Loss | 67% | 0.120 |
| 9 | Tomova et al., 2020 | 29 | Reward | Reward vs. No reward | 38% | 0.929 |
| 10 | Fareri et al., 2012b | 20 | Reward | Social reward  vs.  Nonsocial reward | 55% | 0.409 |
| 11 | Xue et al., 2008 | 19 | Cognitive control | Inhibition success  vs.  inhibition failure | 58% | 0.322 |
| 12 | Wakeman et al., 2015 | 16 | Social cognition | Familiar faces  vs.  unfamiliar faces | 63% | 0.230 |
| 13 | Chen et al., 2019 | 36 | Social cognition | Social scenes  vs.  nonsocial scenes | 47% | 0.685 |
| 14 | Chen et al., 2015 | 27 | Social cognition | Self-referential  vs.  nonself-referential | 40% | 0.873 |
| 15 | Barch et al., 2013 | 484 | Social cognition | Theory of Mind (ToM)  vs.  No ToM | 52% | 0.197 |

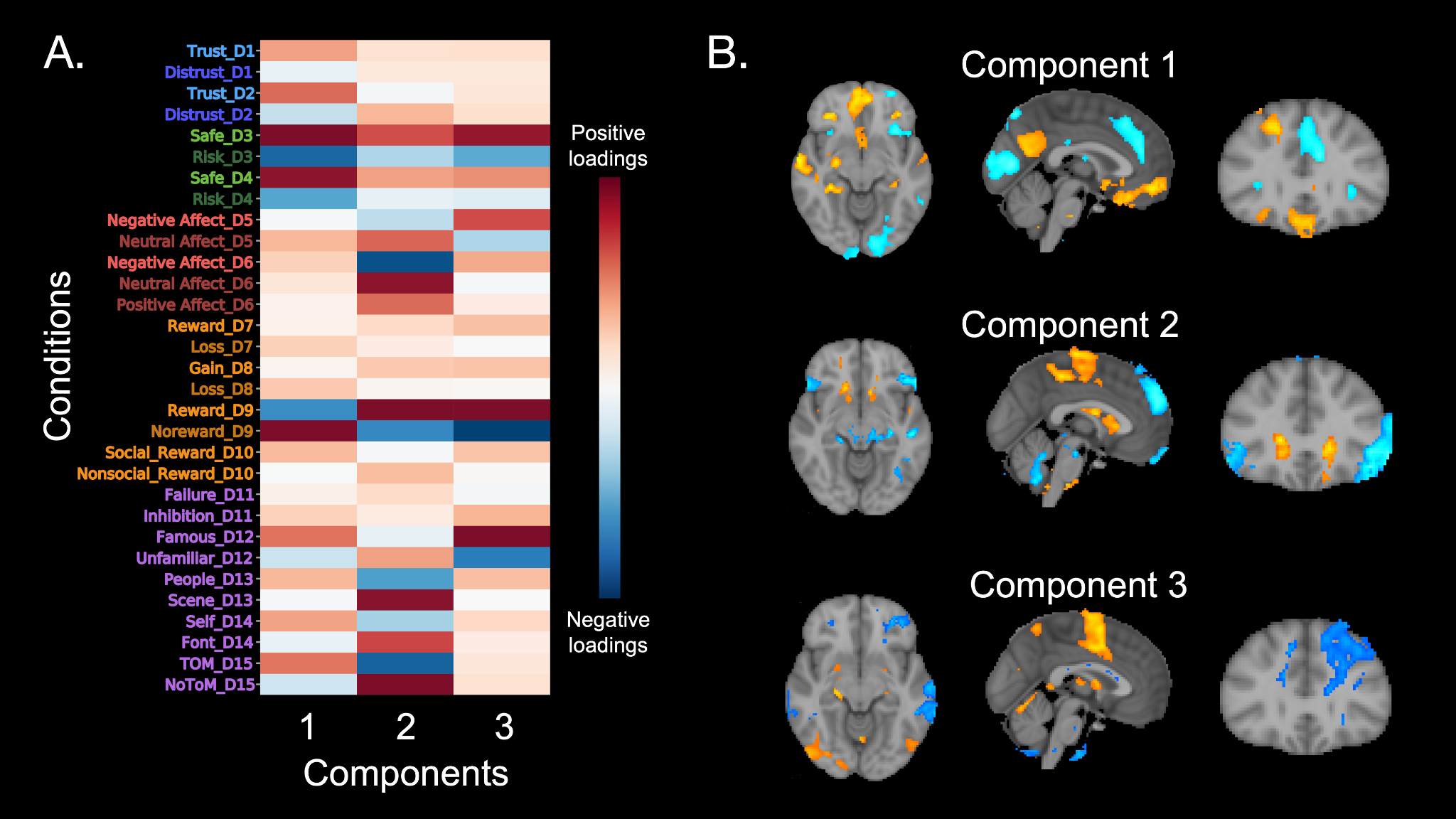

**Figure S1. Condition loadings on each of the three ICA components.** (A) The two trust conditions, safe conditions, no-reward/loss conditions, and one neutral affect condition loaded positively on the first component. By contrast, the two distrust conditions, risk conditions, reward conditions, and one negative affect condition loaded negatively on the first component. (B) Spatial patterns for each of the three components.
